# Long-term hippocampal low-frequency stimulation alleviates focal seizures, memory deficits and synaptic pathology in epileptic mice

**DOI:** 10.1101/2025.01.16.633322

**Authors:** Piret Kleis, Enya Paschen, Andrea Djie-Maletz, Andreas Vlachos, Carola A. Haas, Ute Häussler

**Affiliations:** Experimental Epilepsy Research, Department of Neurosurgery, Medical Center - University of Freiburg, Faculty of Medicine, Freiburg, Germany; Faculty of Biology, University of Freiburg, Freiburg, Germany; Department of Neuroanatomy, Institute of Anatomy and Cell Biology, Faculty of Medicine, University of Freiburg, Freiburg, Germany; BrainLinks-BrainTools Center, University of Freiburg, Freiburg, Germany

**Keywords:** neurostimulation, dentate gyrus, hippocampal sclerosis, kainate, structural plasticity, spatial navigation

## Abstract

**Background:** Mesial temporal lobe epilepsy (MTLE) is a prevalent form of focal epilepsy characterized by seizures originating from the hippocampus and adjacent regions. Neurostimulation presents an alternative for surgery-ineligible patients with intractable seizures. However, conventional approaches have limited efficacy and require refinement for better seizure control. While hippocampal low-frequency stimulation (LFS) has shown promising seizure reduction in animal studies and small clinical cohorts, its mechanisms, sex-specific outcomes, and long-term effects remain unknown.

**Objectives:** We aimed to identify the long-term antiepileptic and cognitive outcomes and potential underlying mechanisms of hippocampal LFS in chronically epileptic male and female mice.

**Methods:** We used the intrahippocampal kainate mouse model replicating the features of MTLE: spontaneous seizures, hippocampal sclerosis, and memory deficits. We applied 1 Hz electrical LFS in the sclerotic hippocampus 6 hours a day, four times a week for 5 weeks and examined its effects on epileptiform activity, spatial memory, and kainate-induced pathological features at cellular and synaptic levels.

**Results:** Long-term hippocampal LFS consistently diminished focal seizures in epileptic male and female mice, with seizure reduction extending beyond the stimulation period. Additionally, LFS relieved spatial memory deficits and reversed pathological long-term potentiation-like changes at perforant path-dentate granule cell synapses. LFS had no significant effect on generalized convulsive seizures, anxiety-like behaviour, neurogenesis, hippocampal sclerosis, excitatory synapse marker expression, or presynaptic vesicles in perforant path fibers.

**Conclusion:** These findings provide clinically relevant insights into the seizure type-specific effects of hippocampal LFS, which, alongside synaptic and behavioural improvements, could contribute to enhanced seizure control and quality of life in MTLE patients.

## Introduction

Mesial temporal lobe epilepsy (MTLE) is the most common form of focal intractable epilepsy in adults, associated with spontaneous recurrent seizures, hippocampal sclerosis, and long-term memory deficits (Engel J 2001; Hoppe et al. 2007; Blümcke et al. 2013). Approximately one-third of epilepsy patients are resistant to antiseizure medications, making invasive treatments such as resective surgery, ablative procedures, or neurostimulation necessary (Semah et al. 1998; Ramey et al. 2013; Kalilani et al. 2018; Ryvlin et al. 2021). Although surgical treatment is the only curative option for pharmacoresistant epilepsy patients, it is not feasible when the seizure focus is not clearly identifiable, lies in the eloquent cortex or several brain regions (Jobst and Cascino 2015; Fois et al. 2016). Neurostimulation via implanted devices offers an alternative for patients ineligible or reluctant to undergo surgery, with the most common approaches being vagus nerve stimulation, deep brain stimulation (DBS) of the anterior nucleus of the thalamus, and closed-loop responsive neurostimulation (RNS) in the epileptogenic zone (Ryvlin et al. 2021). These stimulation approaches are palliative, providing seizure reduction (30-73%) but rarely seizure freedom (0-13%) (Morris and Mueller 1999; Nair et al. 2020; Ryvlin et al. 2021; Salanova et al. 2021; Peltola et al. 2023).

Current neurostimulation strategies can be improved by adjusting targets, parameters, and stimulation modes (Lozano et al. 2019; Mohan et al. 2020). Conventionally, DBS and RNS are applied at high frequencies (HFS, >100 Hz). However, recent small-cohort clinical studies have shown promising seizure reduction with low-frequency stimulation (LFS, <10 Hz) of the hippocampus (Koubeissi et al. 2013; Lim et al. 2016; Koubeissi et al. 2022; Alcala-Zermeno et al. 2023), which is commonly the epileptogenic zone in MTLE (Bartolomei et al. 2005; Tatum 2012). Hippocampal LFS may be more advantageous than HFS due to its lower propensity to induce generalized seizures (Mariani et al. 2021; Sivaraju et al. 2024). Another factor to consider when selecting stimulation settings is the effect on cognitive function. HFS of the entorhinal-hippocampal circuit has demonstrated disruption (Boëx et al. 2011; Nair et al. 2020), enhancement (Nair et al. 2020), or no effect (Velasco et al. 2007; McLachlan et al. 2010; Boëx et al. 2011; Wang et al. 2021) on memory processes, depending on the exact stimulation location, stimulation timing, memory task, brain state, and individual differences. Little is known about the impact of hippocampal LFS on memory in humans (Koubeissi et al. 2013; Koubeissi et al. 2022). In rodent models of MTLE, open-loop and closed-loop hippocampal LFS have been shown to alleviate seizures and spatial memory impairments (Rashid et al. 2012; Salam et al. 2016; Paschen et al. 2020; Ruan et al. 2020; Kleis et al. 2024; Paschen et al. 2024; Zare et al. 2024). However, LFS mechanisms and long-term effects on specific seizure types, hippocampal function, and structural changes have not been investigated. In addition, most studies were performed in males, disregarding possible sex differences.

In the current study, we investigated the effects of repetitive hippocampal LFS in the well-established intrahippocampal kainate (KA) mouse model, which accurately mimics the key pathological features of MTLE, including frequent focal seizures, hippocampal sclerosis, drug resistance, and memory impairment (Bouilleret et al. 1999; Riban et al. 2002; Häussler et al. 2012; Duveau et al. 2016; Van Den Herrewegen et al. 2018; Janz et al. 2018; Masala et al. 2023; Paschen et al. 2024). Like MTLE patients, KA-injected mice experience generalized convulsive seizures infrequently (French et al. 1993; Riban et al. 2002; Sheybani et al. 2018; Kim et al. 2020), necessitating long-term protocols to assess the effects on this seizure type. We applied 1 Hz electrical LFS in the sclerotic hippocampus 6 hours per day, four times a week for 5 weeks, and examined its effect on epileptiform activity, behaviour, and KA-induced pathological features at the cellular and synaptic levels. The optimal stimulus target (dentate gyrus in the sclerotic hippocampus), frequency (1 Hz), and mode (continuous vs. intermittent or on-demand) of LFS were identified in our previous studies (Paschen et al. 2020; Kleis et al. 2024; Paschen et al. 2024). In this preclinical study, we uncover lasting positive effects of hippocampal LFS on focal seizure occurrence and memory performance and reveal synaptic mechanisms potentially underlying these advantageous outcomes.

## Methods

### Study design

Mice were randomly assigned to a saline (Sal, healthy control), KA (epileptic non-stimulated), or KA+LFS (epileptic stimulated) group. Each mouse was subjected to intrahippocampal injections, electrode implantations, electrophysiological recordings, behavioural tests, and histological analysis (Fig. S1). Brain tissue from a subset of mice was processed for electron microscopy and ultrastructural analysis.

### Animals

Experiments were performed in adult transgenic C57BL/6 (Thy1-eGFP-M line (Feng et al. 2000)) male and female mice, aged 9-17 weeks at the start of the experiment. A total of 76 mice bred at the Center for Experimental Models and Transgenic Service in Freiburg were used for this study. Mice were maintained under a 12 h light/dark cycle at room temperature with food and water *ad libitum*. Animals were group-housed before and individually after implantations. All animal experiments complied with EU Directive 2010/63 and were approved by the regional council (Regierungspräsidium Freiburg).

### Kainate injections

Mice were injected with 100 nl KA (5 mM, Tocris) or Sal into the right dorsal hippocampus, as described previously (Janz et al. 2017b; Paschen et al. 2020; Kleis et al. 2024). Mice were deeply anesthetized (ketamine hydrochloride 100 mg/kg (CEVA animal health), xylazine 5 mg/kg (Rompun®, Bayer), atropine 0.1 mg/kg (Braun), i.p.), received analgesic treatment (carprofen 4 mg/kg (Carprieve®, CP-Pharma), s.c.), and placed into a stereotaxic frame (David Kopf Instruments, CA) in flat skull position. The KA or Sal was stereotaxically injected using a Nanoject III (Drummond Scientific Company) at coordinates (in mm): anterioposterior (AP) −2.0, mediolateral (ML) −1.5 relative to bregma, and dorsoventral (DV) −1.5 relative to the cortical surface.

Following intrahippocampal KA injection, status epilepticus was confirmed by observing rotations, immobility, or convulsions, as previously described (Riban et al. 2002; Tulke et al. 2019). The status epilepticus self-terminated within 3-9 hours. Table S1 summarizes the sample sizes and reasons for exclusion.

### Recording and stimulation electrode implantations

Teflon-coated platinum-iridium wires (125 µm diameter; World Precision Instruments) for local field potential (LFP) recordings were implanted 14-17 days after KA/Sal injections into the ipsilateral and contralateral hippocampus (iHC and cHC) at the level of KA injections. Additionally, a stimulation electrode with nanostructured platinum coating (Boehler et al. 2020) was implanted into the ipsilateral dentate gyrus, adjacent to the iHC electrode at a 30° angle (Fig. S1). Stereotaxic coordinates of the recording electrodes were (in mm) AP −2.0, ML +1.4 (cHC), −1.4 (iHC) or −2.4 (stimulation electrode) relative to bregma, and DV −1.6 relative to the cortical surface.

Two stainless steel screws (DIN 84) were implanted above the frontal cortex as reference and ground. Electrodes and screws were soldered to a micro-connector (BLR1-type) and secured with dental cement (Paladur). The electrode positions were verified through post hoc histology as described previously (Paschen et al. 2020).

### Electrophysiological recordings and electrical stimulation

LFPs were recorded in freely moving mice with a tethered system in the chronic phase of epilepsy between 23-57 days after intrahippocampal injections. 6-h recordings were performed in the home cage without a lid, enabling video monitoring. Mice were connected to a miniature preamplifier (MPA8i, Smart Ephys/Multi Channel Systems) and signals were amplified 1000-fold, bandpass-filtered from 1 Hz to 5 kHz and digitized with a sampling rate of 10 kHz (Power1401 analogue-to-digital converter, Spike2 software, Cambridge Electronic Design).

Electrical stimulation was conducted using the 4-Channel stimulus generator STG1004 and MC_stimulus II software (Multi-Channel Systems). Biphasic rectangular current pulses with 400 μs phase duration, anodic first, were applied at 1 Hz. An optimal stimulation current was selected for each mouse in a test recording, where pulses were ramped up stepwise from 15 µA to 200 µA. The optimal current was one that generated a stable maximum response with a minimum current ranging from 100 µA to 200 µA. At the start of each 6-hour stimulation session, a 5-min current ramp (e.g. 20-40-60-80-100 µA) was applied to avoid inducing a generalized seizure as shown in our previous study (Paschen et al. 2024).

### Analysis of epileptiform activity

Hippocampal LFPs were downsampled to 500 Hz and analysed using a semi-automated algorithm that detects and classifies epileptiform activity after manually-controlled removal of artefacts and selection of spike polarity (Heining et al. 2019; Paschen et al. 2020; Heining et al. 2024; Kleis et al. 2024; Paschen et al. 2024). In the intrahippocampal KA mouse model, epileptiform activity occurs during interictal (between-seizure) phases as isolated spikes and spike trains, and during ictal phases as electrographic non-convulsive focal seizures and generalized convulsive seizures (Riban et al. 2002; Twele et al. 2017). The algorithm detects spikes and classifies clusters of spikes, i.e. epileptiform bursts, according to their spike load into low-load (LL), medium-load (ML), and high-load bursts (HL) (Heining et al. 2019; Heining et al. 2024) (Fig. S2a). We considered LL and ML bursts as interictal activity and HL bursts as ictal activity, corresponding to focal seizures (Heining et al. 2019; Kleis et al. 2024). Stimulation-induced responses were disregarded from the classification of epileptiform activity, i.e. the spike detected during an electric pulse was masked. For the focal seizure analysis, we used one reference recording and one recording with LFS per week and calculated the HL burst ratio (time spent in HL burst divided by total recording time). The recording day with LFS was systematically pre-selected to avoid bias: week 1 –Thursday, week 2 – Tuesday, week 3 – Wednesday, week 4 – Friday, week 5 – Monday.

Annotation of generalized seizures was done manually by writing down the start (first discharge of the spike train) and the finish (last spike before post-ictal depression) of the electrographic seizure, and the co-occurrence of the behavioural seizure was confirmed in the video. Generalized seizures and the following 30 min of the LFP recording were excluded from the focal seizure analysis due to post-ictal depression.

### Behavioural experiments

Each mouse was subjected to open-field and light-dark box (LDB) tests twice: once before and once after long-term LFS to assess mobility and anxiety levels (timeline in Fig. S1). On post-KA days 20 and 21, the mice were habituated to the behaviour room and handled each day 2 times for 2 min. The first open-field test (10 min, 35 x 50 cm arena, 60 lx) was conducted on day 22 post-KA and the second open-field test on post-KA day 56. The LDB test was performed on days 23 and 55 post-KA. Mice were placed into the dark compartment and allowed to freely alternate between the dark and light compartments (both 18 × 18 cm, dark: 2 lx and bright: 220 lx) for 10 min.

Barnes maze is a dry-land maze for testing visual spatial learning and memory (Barnes 1979; Pompl et al. 1999). On the elevated (1 m from the ground) circular arena (Ø 1 m), bright illumination (450 lx) encourages the mice to find the escape box, which is below one of the 20 holes, evenly distributed at the edge of the maze. We had eight different spatial cues made of white paper on a black curtain surrounding the arena. During post-KA day 41-44, we habituated the mice to the behavioural test room and handled each mouse for 5 min. On post-KA day 45, the mice were familiarized with the Barnes maze escape box by gently guiding them to enter and letting them stay in the box for 2 min with a chocolate reward. Post-KA days 48-51, each mouse was trained on the Barnes maze 3 times a day with 20-30 min inter-trial interval. For this, the mouse was placed at the centre of the arena under a start box for 10 s, which served as a randomization of the starting direction. The starting box was then lifted from outside the curtain, and the mouse had 3 min to explore the arena before it was guided to the escape box. The mouse had 2 min in the escape box and was then returned to its home cage. 24 hours after the last training, a test trial was performed, where the escape box was removed and the mouse had 90 s on the arena. After each training and test session, the maze was sterilized with 0.5% Incidine and turned 90° to avoid interference of odour cues from previous mice.

During the behavioural tests, animals were video-tracked (Basler acA1300–60 gm overhead camera, 60 Hz, Pylon Camera software) and the videos were analysed using EthoVision XT software (Noldus). We quantified the speed and distance travelled in all tests, time spent in the centre (>7 cm away from the arena wall) in the open-field test, fraction of time spent in the light compartment and number of transitions in the LDB test, time to target and number of errors in Barnes Maze training and test, and search strategy and time spent in quadrants in Barnes maze test. The number of errors was defined as the number of incorrect holes the mouse had been to before its first visit to the target hole. The search strategy was classified as random (randomly crossing the platform before reaching the target location), serial (visiting at least two adjacent holes in series before reaching the target), direct (reaching the target directly or with one adjacent hole next to the target location), or fail (not finding the target).

### Tissue preparation and immunohistochemistry

At the end of each experiment, mice were deeply anesthetized and transcardially perfused with 0.9% saline followed by 4% paraformaldehyde (PFA) in 0.1 M phosphate buffer (PB, pH 7.4). The dissected brains were post-fixed in PFA overnight, transferred to PB, and sectioned (coronal plane, 50 μm) using a vibratome (VT100S, Leica Biosystems). Slices were collected and stored in PB.

We performed immunofluorescence staining for five different markers on separate brain slices: (1) neuronal nuclei (NeuN) and (2) glial fibrillary acidic protein (GFAP) to assess hippocampal sclerosis, (3) doublecortin (DCX) for neurogenesis, (4) vesicular glutamate transporter 1 (vGluT1) and (5) AMPA receptor GluA2 subunit to assess the distribution of excitatory synapses. Free-floating sections were pre-treated with 0.25% (NeuN, GFAP, DCX) or 0.3% (vGluT1, GluA2) TritonX-100 and 10% normal horse serum (Vectorlabs) diluted in PB for 30 min (NeuN, GFAP, DCX) or 60 min (vGluT1, GluA2). Subsequently, slices were incubated with guinea-pig anti-NeuN (1:1000; Synaptic Systems), rabbit anti-GFAP (1:500, Dako), rabbit anti-DCX (1:500, Synaptic Systems), guinea-pig anti-vGluT1 (1:1000, Synaptic Systems) or rabbit anti-GluA2 (1:1000, Synaptic Systems) overnight at 4°C or at RT (vGluT1, GluA2). Sections were rinsed and then incubated for 2.5 h in donkey anti-guinea pig or donkey anti-rabbit Cy5-conjugated secondary antibody (1:200, Jackson ImmunoResearch Laboratories Inc.) followed by extensive rinsing in PB. Counterstaining was performed with DAPI (4′,6-diamidino-2-phenylindole; 1:10.000, Sigma-Aldrich). The sections were mounted on glass slides and coverslipped with Immu-Mount^TM^ medium (Thermo Shandon Ltd.).

### Image acquisition and histological analysis

Tiled fluorescent images of the brain sections were taken with a Zeiss Axio Imager 2 using a Plan-Apochromat 10x objective with a numerical aperture of 0.45 and Zen Software (Carl Zeiss Microscopy GmbH). For each immunolabelling, we imaged three brain slices (−1.46 to −2.46 mm from bregma) with dorsal hippocampus per mouse. The exposure times were kept constant: NeuN – 3.5 s, GFAP – 0.5 s, DCX – 3 s, vGluT1 – 1.5 s, GluA2 – 5 s. Image analysis in Fiji software(Schindelin et al. 2012) was performed by a blinded investigator.

To assess the severity of hippocampal sclerosis, we quantified granule cell dispersion and neuronal loss in NeuN-labelled sections and astrogliosis in GFAP-labelled sections. We measured the width of the granule cell layer (GCL) by drawing three lines in the upper blade and three lines in the lower blade, perpendicular to the GCL outline. To estimate the extent of cell loss in the Cornu ammonis (CA) regions, we subtracted the remaining CA length (usually CA2) from the theoretical total CA length (from the border with fasciola cinereum to the point in the hilus where CA3 usually finishes). For neuronal loss percentage, we divided the length of lost CA by the theoretical total CA length. In GFAP-labelled sections, we drew a region of interest (ROI) around the ipsilateral hippocampus, excluding the areas with glial scarring around the electrodes, and measured the mean grey value. In each slice, we normalized the mean grey values by division with the mean grey values of the background (150 × 150 µm square in L5-6 in the contralateral primary somatosensory cortex).

In DCX-labelled sections, we counted DCX-positive cells in the contralateral dentate gyrus and measured the GCL length at the inner surface (towards the hilus). DCX-positive cell count per 1 mm GCL was calculated. GluA2 and vGluT1-labeled sections were analysed for fluorescence intensity in five layers of the dentate gyrus: outer molecular layer, middle molecular layer, inner molecular layer, GCL, and hilus. In each layer, we measured the mean grey value in three small ROIs (35 × 35 µm), which were then averaged. The values were normalized to the local background measured in a 128 × 128 µm square in the corpus callosum.

### Electron microscopy and ultrastructural analysis

Mice, in which the brain tissue was used for transmission electron microscopy (males: n=8, Table S1), received a recombinant adeno-associated virus (AAV1.CaMKIIα.hChR2(H134R)- mCherry.WPRE.hGH, 450 nl, physical titre: 6.5 x 10E12 vg/ml, Viral Vector Facility University of Zurich) injection with Nanoject III into the medial entorhinal cortex (coordinates from bregma: ML = −2.9 mm; from transverse sinus: AP = +0.2 mm; from cortex surface: DV = −1.8 mm; injection angle: 9°) during the same surgery as implantations. The injection needle was left at the injection site for five minutes after injection to avoid reflux.

Immediately after the last stimulation/recording on post-KA day 57, mice were deeply anesthetized and transcardially perfused with 4% PFA and 0.1% glutaraldehyde in 0.1 M PB. The tissue was post-fixed for 1 h, stored in PB and sectioned with a vibratome as described above. We selected four brain slices (per mouse) in the vicinity of the stimulation electrode. For immunogold labelling, free-floating sections were pre-incubated for 30 min with 20% normal goat serum, followed by incubation with rabbit anti-mCherry (1:100; Invitrogen) and then with goat anti-rabbit 1.4 nm gold-coupled secondary antibody (1:100, Nanogold; Nanoprobes), both at 4°C overnight. The next day, immunogold labelling was enhanced with a HQ silver kit (Nanoprobes). Sections were then osmicated with 0.5% OsO_4_ in 0.1 M PB for 30 min, stained with 1% uranyl acetate in 70% ethanol for 30 min, dehydrated in graded ethanol, and flat-embedded in epoxy resin (Durcupan ACM; Sigma-Aldrich). Finally, the re-embedded sections were trimmed to the molecular layer and ultrathin sections were cut with a diamond knife.

Transmission electron microscopy was performed using a LEO 906E electron microscope (Carl Zeiss Microscopy GmbH). Photomicrographs from random sections were obtained, and PP- DGC synapses (60 per mouse) were analysed using ImageSP software (Sysprog) by a blinded investigator. PP-DGC synapses were identified by presynaptic nanogold-labelling in the MML of the dentate gyrus. Only synapses with distinct membrane borders, clearly visible presynaptic vesicles, and postsynaptic densities (PSDs) were selected for quantification (50 synapses per mouse). At each selected PP synapse, we counted presynaptic vesicles and measured the area of PP boutons, putative DGC spine heads (30 per mouse), mitochondria within PP boutons, and the length of PSDs. We calculated vesicle density by dividing the presynaptic vesicle count by the mitochondria-free bouton area. The fractions of (1) boutons with multiple active zones, (2) boutons with multiple spines, (3) multiple PSD spines, and (4) perforated synapses characterized by discontinuous PSDs were determined by two independent observers blinded to experimental groups (KA / KA+LFS).

### Statistics

Statistical analyses were performed with GraphPad Prism 10. We used the Shapiro-Wilk test for normality to select an appropriate parametric or non-parametric test. If the data fit a lognormal rather than a normal distribution, we applied statistical tests on log-transformed data. To compare two independent data sets at one time point (or the average of time points), we used either an unpaired t-test for parametric data or the Mann-Whitney test for non-parametric data. Multiple comparisons without the time factor were performed using (1) one-way ANOVA with Šídák’s multiple comparisons test for parametric data, (2) Kruskal-Wallis test with Dunn’s multiple comparisons test for non-parametric data, (3) multiple paired t-tests with correction for parametric data, or (4) Mann-Whitney tests with multiple comparison correction for non-parametric data. To compare groups at different time points, we applied two-way repeated measures (RM) ANOVA with Tukey’s (comparing all groups), Šídák’s (comparing pre-selected pairs of data) or Dunnett’s (comparing values to the baseline) multiple comparisons test. All statistical tests were two-sided. The significance thresholds were set to *P<0.05, **P<0.01, and ***P<0.001 (same for Q-values that incorporate multiple comparisons correction). Data are represented as mean (parametric data) or median (non-parametric data or dataset with n<4) with 95% CI as error bars. Experimenters were blinded to the experimental group (KA vs. KA+LFS) for histological, ultrastructural, and behavioural analyses. Statistical methods were not applied to pre-determine sample sizes, however, our sample sizes are comparable to those used in previous studies (Janz et al. 2017b; Jafari et al. 2021; Chen et al. 2023; Paschen et al. 2024).

## Results

### Persistent reduction of focal seizures by hippocampal LFS

Focal seizures are the most frequently observed seizure type in both MTLE patients and intrahippocampal KA mice (Engel 1995; Bouilleret et al. 1999; Riban et al. 2002; Cendes 2005; Janz et al. 2018; Kim et al. 2020). Here, we compared electrographic focal seizures and their progression in male and female intrahippocampal KA mice that were either recorded and stimulated (KA+LFS) or only recorded (KA) with intrahippocampal electrodes (Fig. 1A-B). All mice were recorded for a 6-h reference LFP once per week and the KA+LFS mice received continuous 6-h-long hippocampal LFS (1 Hz) four times per week for five weeks (Fig. 1C). Using the PEACOC algorithm(Heining et al. 2019; Heining et al. 2024), epileptiform activity in the iHC was detected and classified as LL, ML, and HL bursts of epileptiform activity (Fig. 1D).

**Figure 1.**
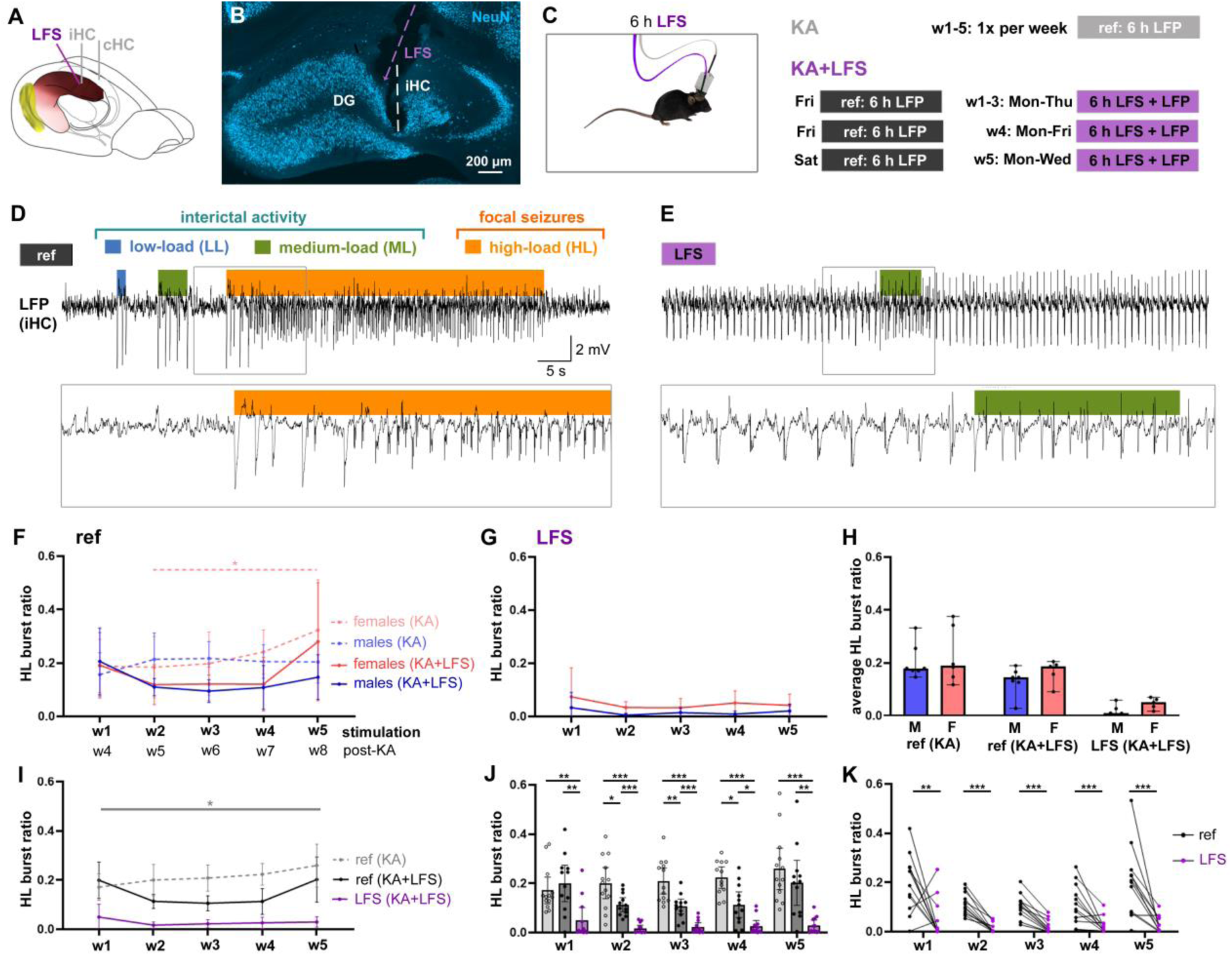
Long-term LFS persistently reduces focal seizures in both sexes. (A) Implantation scheme of recording (ipsilateral hippocampus − iHC, contralateral hippocampus − cHC) and stimulation (LFS) electrodes. (B) Representative image of a NeuN-labelled hippocampal section showing location of the iHC recording and LFS electrode in the dentate gyrus (DG) of the sclerotic hippocampus. (C) Freely moving mice in their home cages were recorded (and stimulated) for 6 hours per day. KA mice (gray) and KA+LFS mice (black) were recorded for reference (ref) once a week, KA+LFS mice received LFS (purple) on 4 days a week. (D) Representative LFP trace from iHC during a ref recording showing interictal activity (LL and ML bursts) and a focal seizure (HL burst). (E) Representative LFP during LFS showing responses to 1 Hz electrical pulses and a ML burst. (F) HL burst ratio in KA females (dashed light red, n=6) and males (dashed light blue, n=7) as well as KA+LFS females (red, n=5) and males (blue, n=7) were comparable during ref recordings across 5 weeks. Two-way RM ANOVA (KA mice: interaction: F_(4, 44)_=1.82, P=0.14, time factor: F _(3,208, 35,29)_=4.086, P=0.012, group factor: F_(1, 11)_=0.167, P=0.69; KA+LFS mice: interaction: F_(4, 40)_=1.19, P=0.33, time factor: F_(2,023, 20,23)_=3.85, P=0.038, group factor: F_(1, 10)_=1.34, P=0.27). Tukey’s multiple comparisons test for time factor. (G) HL burst ratio in KA+LFS females and males during LFS across 5 weeks. Two-way RM ANOVA (interaction: F_(4, 40)_=0.19, P=0.94, time factor: F_(1,311, 13,11)_=1.08, P=0.34, group factor: F_(1, 10)_=6.16, P=0.032), Šídák’s multiple comparisons test between males and females at each week. (H) Average HL burst ratio for all 5 weeks in males and females in each group reveals no sex differences. Mann-Whitney tests with multiple comparison correction (ref KA: Q=0.63; ref KA+LFS: Q=0.31; stim KA+LFS: Q=0.09). (I) Progression of HL burst ratio in five recording weeks in KA (n=13) but not KA+LFS (n=12) mice (both sexes) during ref and LFS. Two-way RM ANOVA (interaction: F_(8, 136)_=2.86, P=0.0057, time factor: F_(2,673, 90,89)_=3.71, P=0.018, group factor: F_(2, 34)_=31.49, P<0.0001) with Tukey’s multiple comparisons test for time factor. (J) HL burst ratio is significantly lower in the ref recordings of KA+LFS mice (filled circles) than in KA mice (unfilled circles) during w2-4. During LFS, HL burst ratio is reduced compared to ref in KA and KA+LFS mice. Two-way RM ANOVA (interaction: F_(8, 136)_=2.86, P=0.0057, time factor: F_(2,673, 90,89)_=3.71, P=0.018, group factor: F_(2, 34)_=31.49, P<0.0001) with Tukey’s multiple comparisons test for group factor. (K) Pairwise comparison of HL burst ratio in ref and LFS recordings of KA+LFS mice. Wilcoxon matched-pairs signed rank test (w1: Q=0.0042, w2-w4: Q=0.00016, w5: Q=0.00086). *P<0.05, **P<0.01, ***P<0.001. Data are presented as mean (f, g, i, j) or median (h) with 95% CI.

The fraction of time spent in focal seizures (i.e. HL burst ratio) during reference recordings was similar between females and males in both KA and KA+LFS groups across five weeks although KA females exhibited an increase from week 5 to week 8 after KA (Fig. 1F). During LFS sessions, the HL burst ratio was consistently low in both sexes with slightly higher values in females than males (Fig. 1G). The average burst ratios during reference recordings in KA and KA+LFS mice, as well as during LFS were not significantly different between males and females (Fig. 1H). Therefore, we pooled the results from both sexes to further investigate the effects of long-term LFS on electrographic epileptiform activity.

Our findings reveal that the HL burst ratio significantly increased over the 5-week period in KA mice but not in KA+LFS mice (Fig. 1I), indicating that long-term LFS might hinder the progression of focal seizures during the chronic phase of MTLE. Before stimulation began in the first week, KA and KA+LFS mice had similar HL burst ratios. However, at weeks 2-4 KA+LFS mice had significantly lower HL burst ratios in the reference recordings (Fig. 1J), suggesting that long-term LFS has a seizure-suppressive effect lasting at least 24 h.

During LFS, the HL burst ratio was significantly reduced compared to the reference recordings of KA or KA+LFS mice in the same week (Fig. 1J). In addition, a pair-wise comparison of the HL burst ratio in the reference and LFS recordings in the same mice confirmed a consistent reduction in focal seizures by hippocampal LFS in all five weeks (Fig. 1K). The LFS-mediated reduction in HL burst ratio resulted from both, fewer burst numbers and shorter burst durations (Fig. S2A-C). Furthermore, hippocampal LFS shortened ML bursts and reduced the number and duration of LL bursts, indicating a positive effect on interictal activity (Fig. S2D-G). Together, these results demonstrate that hippocampal LFS can consistently reduce focal seizures in chronically epileptic male and female mice over long periods, with a strong transient seizure-suppressive effect during LFS and a weaker lasting effect at least 24 hours after LFS.

### Hippocampal LFS has no effect on generalized convulsive seizures

Although generalized convulsive seizures are rare in patients with MTLE and intrahippocampal KA mice, they are clinically significant due to their severity and increased risk for sudden, unexpected death (Engel 1995; Bouilleret et al. 1999; Duveau et al. 2016; Sheybani et al. 2018). Thus, long-term stimulation and recording are required to assess the effects of hippocampal LFS on spontaneous generalized seizures. We counted the generalized seizures with behavioral manifestations in every recording with and without LFS (Fig. 2A, B). We report that generalized seizures in KA+LFS mice occurred with similar frequency in recordings with and without LFS (Fig. 2C). A comparison of the generalized seizure numbers in KA and KA+LFS mice revealed no difference in male mice (Fig. 2D). Surprisingly, three of six KA+LFS females had a high generalized seizure occurrence, similar to mice with strong hippocampal atrophy (4 males, 2 females, Fig. S3). Female KA+LFS mice and mice with atrophy experienced slight but not statistically significant progression in the generalized seizure number over weeks (Fig. 2E). Generalized seizure duration was not altered in KA+LFS mice compared to that in KA mice (Fig. 2F). Overall, long-term hippocampal LFS did not reduce the incidence or duration of generalized convulsive seizures.

**Figure 2.**
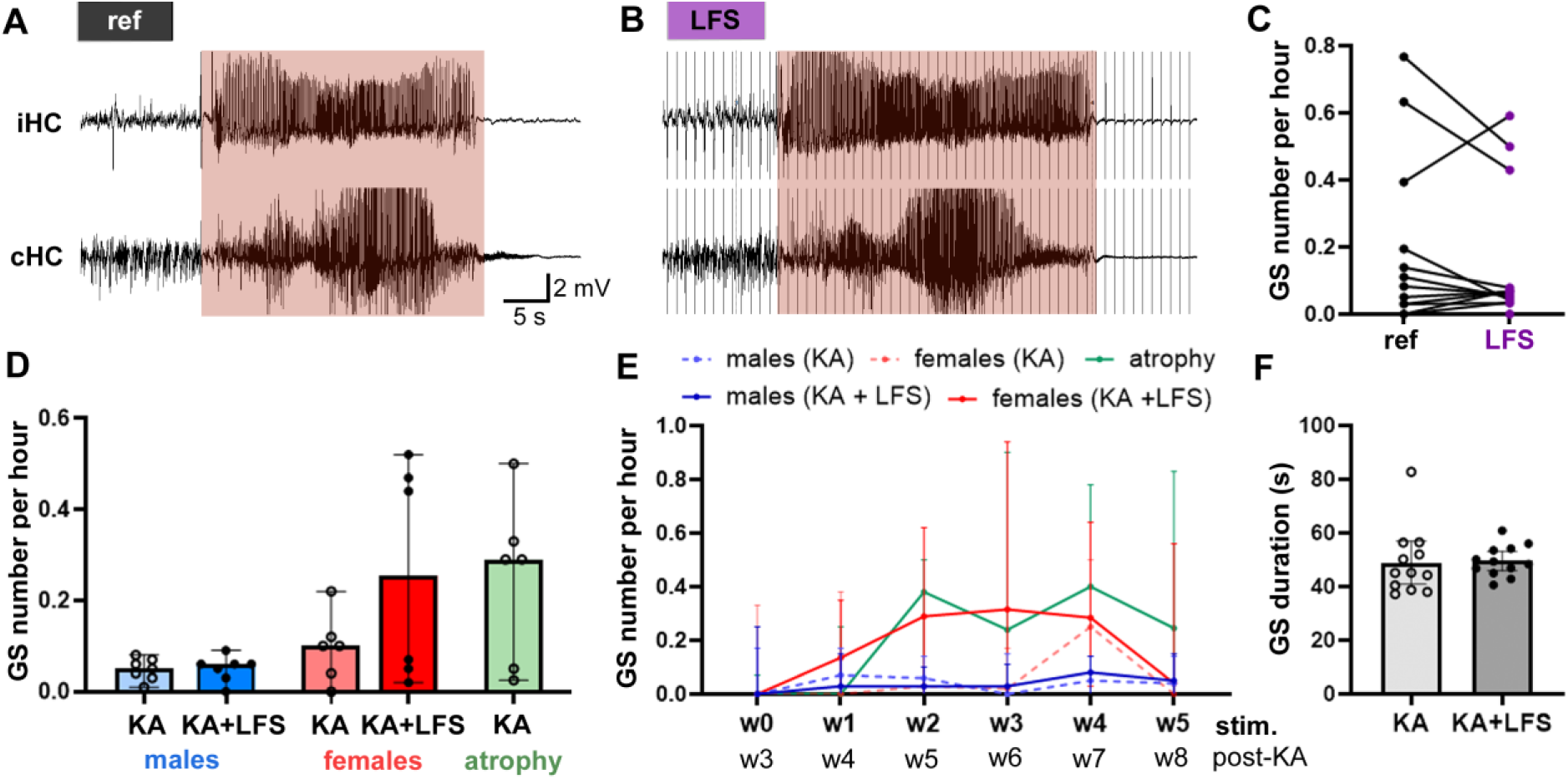
Hippocampal LFS has no effect on generalized convulsive seizures. (A, B) Representative generalized seizure (GS) during ref (A) and LFS (B) in the same mouse. (C) GS number per hour during ref and LFS recordings (n=12). Paired t-test (T_12_=0.94, P=0.37, n=13). (D) GS number per hour in male KA (n=7) and male KA+LFS (n=7) mice, female KA (n=6) and female KA+LFS (n=6) mice and in KA mice with severe hippocampal atrophy (n=6). Unfilled circles – unstimulated mice, filled circles – stimulated mice. Kruskal-Wallis test (P=0.24). (E) Progression of GS number from baseline (w0) to the fifth stimulation/recording week (w5). Mixed-effects analysis (interaction: F_(20, 132)_=2.41, P= 0.0016, time factor: F_(2,975, 78,55)_=6.69, P=0.0005, group factor: F_(4, 27)_=3.83, P=0.014), Dunnett’s multiple comparisons test. (F) GS duration in KA (n=12) and KA+LFS (n=12) mice. Unpaired t-test (T_22_=0.16, P=0.87). Data are presented as median (G-E) or mean (F) with 95% CI.

### Long-term LFS improves spatial memory deficits in epileptic mice

Determining the long-term effects of LFS on hippocampal function is important, particularly given the association between MTLE and memory impairment (Helmstaedter et al. 2003; Tramoni-Negre et al. 2017; Van Den Herrewegen et al. 2018). To assess spatial learning and memory recall, we employed the Barnes maze test (see Methods) in healthy, epileptic, and stimulated epileptic mice. The Barnes maze was conducted seven weeks after intrahippocampal injections (Sal/KA) in the fourth stimulation week (Fig. 3A, see Methods). We found that some epileptic female mice (n=5) and mice with pronounced hippocampal atrophy (n=5) which exhibited a high generalized seizure rate during the week of Barnes maze training showed freezing behavior at the center of the arena or jumped off the maze (Fig. 3B). These mice were excluded from the behavioral analysis.

**Figure 3.**
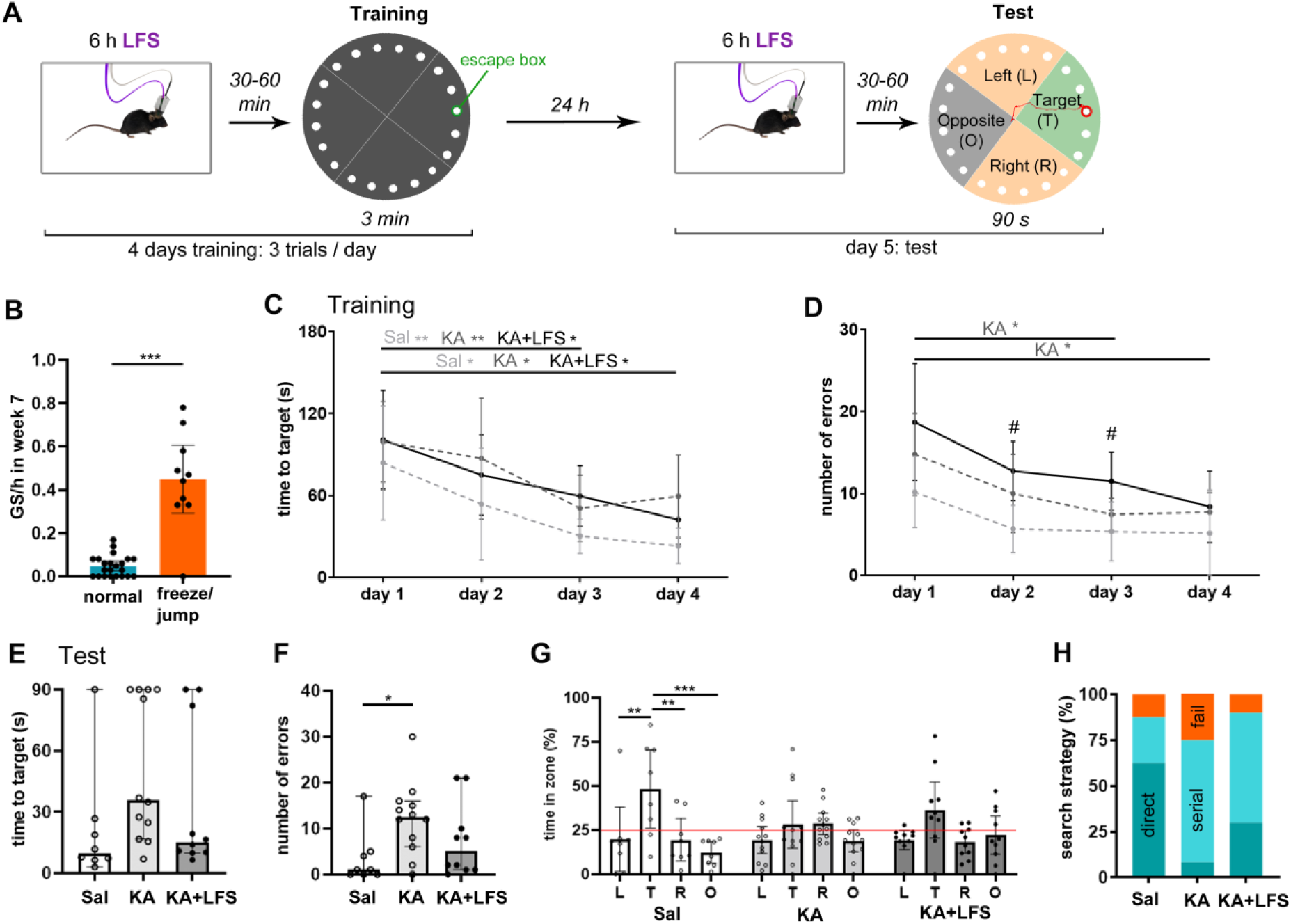
Long-term LFS improves spatial learning and memory in epileptic mice. (A) Spatial learning and memory were assessed in Barnes maze training (escape box under the target hole, in green) and test (escape box removed, in red). (B) Some mice were excluded from the Barnes maze test due to freezing or jumping off the maze. These mice (n=10), had significantly higher generalized seizure (GS) occurrence during the week of Barnes maze training (week 7) compared to mice that could perform the test (n=22). Mann Whitney test. (C) All mice, including Sal (n=8: 5 males, 3 females; light gray intermittent line), KA (n=12: 7 males, 5 females; dark gray intermittent line), and KA+LFS (n=10: 7 males, 3 females; black line) mice got significantly faster in finding the target over 4 training days. Two-way ANOVA on log-transformed data (interaction: F_(6, 81)_=0.73, P=0.63, time factor: F_(2,322, 62,71)_=18.67, P<0.0001, group factor: F_(2, 27)_=2.49, P=0.10) with Tukey’s multiple comparison test across days. (D) The number of errors dropped significantly in KA mice across 3 and 4 days. Two-way ANOVA (interaction: F_(6, 81) =_ 0.47, P=0.96, time factor: F_(2,777, 74,99)_=10.28, P=0.001, group factor: F_(2, 27)_= 6.96, P= 0.0037) with Tukey’s multiple comparison test across days (*) and between groups (difference between Sal and KA+LFS: #P<0.05). (E) Time to target during the test trial. Kruskal-Wallis test (P=0.056). (F) Number of errors in the test trial was significantly higher in KA mice than in Sal mice. Kruskal-Wallis test (P=0.038), Dunn’s multiple comparisons test. (G) Quadrant preference during the test trial, red line indicates the chance level (25%). Two-way ANOVA (interaction: F_(6, 108)_=2.08, P=0.061, quadrant: F_(3, 108)_=9.81, P<0.0001, group factor: F_(2, 108)_= 0.063, P=0.94) with Tukey’s multiple comparisons test. (H) Fraction of search strategies (direct: dark turquoise, serial: light turquoise) and failures (fail: orange) in finding the target in the test trial. *P<0.05, **P<0.01, ***P<0.001. Data are presented as mean (B-D, G) or median (E, F) with 95% CI.

In the training phase of the Barnes maze, mice in all groups got faster in reaching the target: the average time to target on day 3 and 4 was significantly lower than on day 1 (Fig. 3C). The number of errors also decreased over 4 days in all groups, however, only KA mice demonstrated statistically significant improvement on day 3 and 4 compared to day 1 (Fig. 3D). Group-wise comparisons of error numbers revealed significant differences between Sal and KA+LFS mice on day 2 and 3 but not on the last training day. These results suggest that both unstimulated and stimulated epileptic mice can learn the Barnes maze task unless they have a high amount of generalized seizures.

We investigated long-term memory in the test trial, 24 h after the last training, where the escape box was removed (Fig. 3A). KA mice tended to require more time finding the target, often not reaching it in 90 seconds (4 out of 12 mice; Fig. 3E), and made significantly more errors than Sal mice (Fig. 3F). Interestingly, KA+LFS mice did not perform significantly different from Sal mice in terms of time to target and number of errors. In the test, Sal mice spent considerably more time in the target quadrant than in the other quadrants, whereas KA and KA+LFS mice did not have a significant preference (Fig. 3G). Nevertheless, most KA+LFS mice (70%) spent more time in the target quadrant than chance level, suggesting an improved memory compared to KA mice (50% above chance level). Most of the Sal mice (62.5%) went to the target directly, whereas KA (60%) and KA+LFS (67%) preferentially used a serial search strategy to find the target in the Barnes maze test (see Methods, Fig. 3H). Importantly, KA+LFS mice were more likely to use a direct search (30%) and less likely to fail in finding the target (10%) than KA mice (direct: 8%, fail: 25%).

Additionally, we analyzed mobility (distance and velocity) and anxiety-like behavior in the open-field and light-dark box tests, where LFS-induced changes were not apparent (Fig. S4).

However, KA mice spent significantly less time in the center of the open-field arena than Sal mice (Fig. S4A, B), suggesting anxiety or thigmotaxis, the tendency to stay near the perimeter. Collectively, these data indicate that epileptic mice use a non-spatial strategy to perform in the Barnes maze, potentially due to spatial memory impairment, which is slightly improved by long-term LFS.

### Extended LFS does not alter hippocampal sclerosis or neurogenesis

Next, we compared the extent of cell loss, astrogliosis, and neurogenesis in the hippocampi of unstimulated and stimulated epileptic mice from both sexes. We quantified the GCL width to characterize the degree of granule cell dispersion and estimated the percentage of neuron loss in the CA areas in NeuN-labeled hippocampal sections (Fig. 4A, B). GCL width and cell loss in CA regions were similar in the KA and KA+LFS mice in both sexes (Fig. 4C, D). In the sclerotic ipsilateral and the contralateral hippocampus, the intensities of GFAP labeling, an indicator of astrogliosis, were higher in epileptic mice than Sal but not different between KA and KA+LFS male and female mice (Fig. 4E-H). Notably, contralateral astrogliosis was the only histological parameter correlating with generalized seizure activity (Fig. S5). All in all, the extent of hippocampal sclerosis was similar in unstimulated and stimulated epileptic mice.

**Figure 4.**
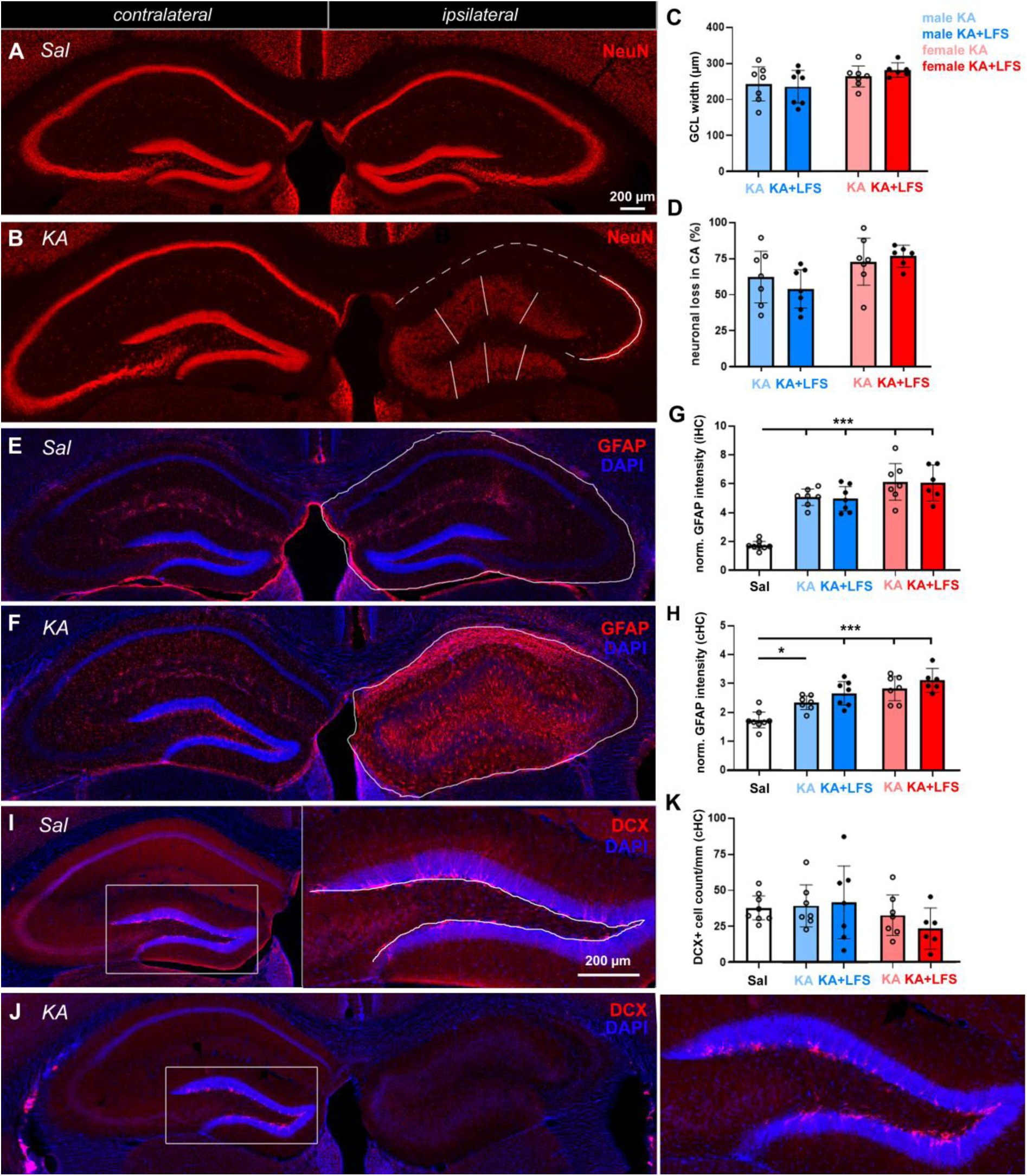
Lack of LFS-induced effects on hippocampal sclerosis and neurogenesis. (A) Representative NeuN-labeled hippocampal section from a Sal-injected mouse. (B) Representative NeuN-labeled hippocampal section from epileptic kainate (KA)-injected mouse. Solid lines in the DG indicate the positions where GCL width was measured (3 sections/mouse). Neuronal loss in CA regions was estimated by measuring the surviving CA length (solid line) and the predicted lost CA length (dashed line). (C) Comparison of GCL width in male KA (light blue, n=7) and KA+LFS (blue, n=7) mice and female KA (light red, n=7) and KA+LFS mice (red, n=6). One-way ANOVA (F(3, 23)=1.66, P=0.20). (D) Loss of neurons in CA regions (percentage of cell loss from total CA length) in male and female KA and KA+LFS mice. One-way ANOVA (F(3, 23)=2.89, P=0.057). (E-F) GFAP staining in (E) Sal and (F) KA mice, showing strong astrogliosis in the ipsilateral hippocampus (iHC). The area of GFAP intensity measurement is marked with a solid line. (G, H) Normalized (norm.) GFAP intensity in Sal, KA and KA+LFS mice in (G) ipsilateral and (H) contralateral hippocampus (cHC). One-way ANOVA (iHC: F(4, 30)=27.35, P<0.0001, cHC: F(4, 30)=14.33, P<0.0001), Sidak’s multiple comparisons test (pairs: Sal with every other group, male KA and KA+LFS, female KA and KA+LFS, females and males with KA, females and males with KA+LFS), *P<0.05, ***P<0.001. (I, J) DCX-labeled hippocampal sections for assessment of neurogenesis in (I) Sal (contralateral) and (J) KA mice (contralateral and ipsilateral). (K) DCX-positive cell counts per 1 mm in the contralateral hippocampus of Sal, KA and KA+LFS mice. One-way ANOVA (F(4, 30)= 1.15, P=0.35). *P<0.05, ***P<0.001. Data are presented as mean with 95% CI.

We hypothesized that LFS could impact epileptiform activity and spatial memory by modulating neurogenesis that is disturbed in chronically epileptic mice (Heinrich et al. 2006; Ledergerber et al. 2006; Häussler et al. 2012; Moura et al. 2020). Accordingly, we did not find any DCX-positive newly generated neurons in the sclerotic hippocampus of KA mice at 8 weeks after KA (Fig. 4J). However, LFS did not modify neurogenesis in the ipsilateral hippocampus, where the neurogenic niche is largely depleted (Sierra et al. 2015), nor in the contralateral hippocampus (Fig. 4I-K). These results demonstrate that long-term hippocampal LFS has no effect on neurogenesis in epileptic mice.

### LFS reverses LTP-like ultrastructural changes in the dentate gyrus

Previously, it was found that intrahippocampal KA mice exhibit LTP-like ultrastructural changes at PP-DGC synapses (Janz et al. 2017a). These changes include enlarged PP boutons and DGC spines, an increased proportion of spines with multiple postsynaptic densities (PSDs), spinule (postsynaptic processes that invade the presynaptic terminal) synapses, and multiple spine boutons. Here, we investigated whether the mechanisms underlying LFS-induced seizure suppression involve changes in synaptic plasticity, such as vesicle depletion or structural rearrangements at the presynaptic and postsynaptic compartments of the PP-DGC synapse. To this end, we injected a viral vector carrying the fluorescent marker mCherry under the control of calmodulin-dependent protein kinase IIα (CaMKIIα) promoter into the medial entorhinal cortex to trace PP fibers in a subset of KA and KA+LFS mice (see Table S1). We then performed immunogold labeling of mCherry positive fibers and transmission electron microscopy to identify PP boutons and characterize PP-DGC synapses in the MML, close to the LFS-stimulation site (Fig. 5A-C, Fig. S1). We found that presynaptic vesicle density at the labeled PP boutons was not altered in KA+LFS mice (n=3, 50 synapses per mouse) compared to KA mice (n=3, 50 synapses per mouse, Fig. 5E-F), suggesting that vesicle pool depletion was not the seizure-suppressive mechanism of LFS.

**Figure 5.**
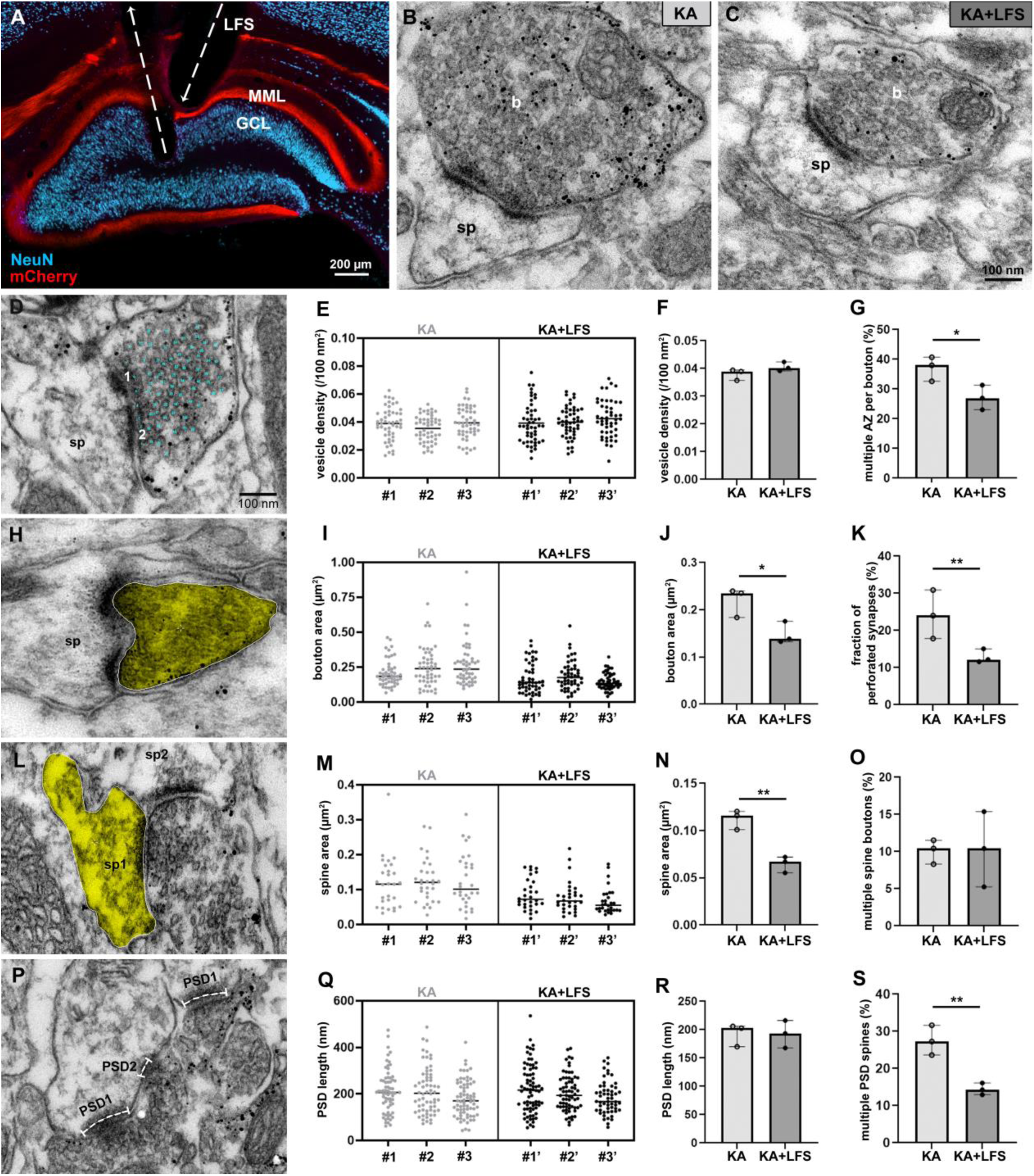
LFS reverses LTP-like ultrastructural changes in the dentate gyrus. (A) Tracing of PP fibers (mCherry, in red) in the MML of the sclerotic hippocampus. (B, C) Representative EM images of nanogold-labeled mCherry-positive PP boutons (b) and putative dentate granule cell spines (sp) in (B) a KA and (C) a KA+LFS mouse. (D) Quantification of presynaptic vesicles (cyan dots) and number of active zones (AZ: 1,2). € Vesicle density in 50 PP boutons per mouse, individual values with mean (normal distribution). (F) Mean vesicle density in each KA (n=3) and KA+LFS (n=3) mouse. Unpaired t-test (T_4_=1.70, P=0.17). (G) Fraction of boutons with multiple AZs was significantly lower in KA+LFS than KA mice. Unpaired t-test (T_4_=2.96, P=0.041). (H) Example of a perforated synapse and quantification of the bouton area (yellow). (I) Bouton area in 50 synapses per mouse, individual values with median (lognormal distribution). (J) Bouton area in KA+LFS mice was significantly lower than in KA mice. Unpaired t-test (T_4_=3.15, P=0.035). (K) Fraction of perforated synapses was significantly reduced in KA+LFS mice compared to KA mice. Unpaired t-test (T_4_=2.88, P=0.035). (L) Quantification of putative dentate granule cell spine area (yellow) and spine number per PP bouton. (M) Spine area in 30 spines per mouse, individual values with median (lognormal distribution). (N) Spine area (median) was lower in KA+LFS compared to KA mice. Unpaired t-test (T_4_=6.20, P=0.0034). (O) Fraction of boutons with multiple spines. Unpaired t-test (T_4_=0.084, P=0.94). (P) Quantification of PSD length and number (PSD1, PSD2). (Q) PSD length in 50 synapses, individual values with mean (normal distribution). (R) Mean PSD length in each KA and KA+LFS mouse. s, Fraction of spines with multiple PSDs. Unpaired t-test (T_4_=5.23, P=0.0064). *P<0.05, **P<0.01. Data are presented as median with 95% CI.

Regarding ultrastructural modifications at the PP-DGC synapse, KA+LFS mice had significantly reduced (1) fraction of boutons with multiple active zones (Fig. 5G), (2) bouton area (Fig. 5H-J), (3) fraction of perforated synapses (Fig. 5H, K), (4) spine area (Fig. 5L-N), and (5) fraction of spines with multiple PSDs (Fig. 5P, S). The fraction of multiple spine boutons (Fig. 5L, O), PSD length (Fig. 5P, R), and the expression of presynaptic and postsynaptic markers of excitatory synapses at the microscopic level were not altered by long-term LFS (Fig. S6). To sum up, LFS reversed several LTP-like pathological changes at the PP-DGC synapse without altering the overall quantity of excitatory synapses.

## Discussion

In the present study, we found that long-term hippocampal LFS persistently suppressed focal seizures in chronically epileptic male and female mice, accompanied by improvements in spatial memory and ultrastructural remodelling at the PP-DGC synapse. Extended recording and stimulation times allowed us to assess the impact of LFS on generalized seizure occurrence, which remained unaffected. Additionally, LFS did not induce changes in hippocampal sclerosis, neurogenesis, expression of excitatory synapse markers, or presynaptic vesicle density in PP boutons near the stimulation site.

### Lasting suppression of focal seizures by LFS: time course and possible mechanisms

In addition to the reliable suppression of focal seizures during continuous 1 Hz stimulation at the sclerotic hippocampus, we observed a reduction in focal seizures during the stimulation OFF phases, at least 24 h after the last stimulation session. Previous studies have also reported long-term seizure reduction in rat models of MTLE following intermittent or closed-loop hippocampal LFS, with effects lasting up to two weeks (Rashid et al. 2012; Zare et al. 2024). In our study, repeated LFS impeded the progression of focal seizures during the second month after intrahippocampal KA injection. Other studies have reported no change in the focal seizure number over one to two months in male mice (Riban et al. 2002; Pallud et al. 2011) or both sexes pooled (Lisgaras and Scharfman 2022) and a significant increase during two months in female mice (Widmann et al. 2024), confirmed by our data in non-stimulated chronically epileptic mice.

The seizure reduction during stimulation OFF phases may be linked to LFS-induced synaptic changes. Intrahippocampal KA mice show LTP-like changes, including enlarged, perforated, and multiple spine PP-DGC synapses, which could drive seizure activity in the sclerotic hippocampus (Janz et al. 2017b). LFS is known to induce long-term depression (LTD) at hippocampal pyramidal synapses, though LTD at PP-DGC synapses is difficult to elicit in slices from healthy (Abraham et al. 1996; Doyère et al. 1996; Fung et al. 2011; Gonzalez et al. 2014) and epileptic rodents (Bragin et al. 2002; Toprani and Durand 2013) or humans (Beck et al. 2000). In the current study, we showed that LFS induced LTD-like structural modifications (Nägerl et al. 2004; Becker et al. 2008), such as a reduction in PP bouton and DGC spine size and fewer perforated synapses. Spine shrinkage, potentially linked to homeostatic downscaling, could also explain these alterations (Wang et al. 2007). Given that associative and homeostatic plasticity last from hours to days (Manahan-Vaughan and Braunewell 1999; Hobbiss et al. 2018), these forms of plasticity could account for the persistent focal seizure reduction. In fact, clinical studies have shown that seizure control by RNS relies on plastic changes and functional reorganization of networks, rather than the immediate effects of seizure interference by closed-loop stimulation (Kokkinos et al. 2019; Khambhati et al. 2021; Rao and Rolston 2023; Anderson et al. 2024). Importantly, we sacrificed the mice within 10 min after the last LFS session, preventing us from determining whether the ultrastructural changes result from short-term or long-term LFS effects. We aimed to test if presynaptic vesicle depletion in PP boutons accounts for the previously known seizure suppression during and 10 min after LFS (Paschen et al. 2020; Paschen et al. 2024), reasoning the short time interval between the stimulation and sacrifice. Our results suggest that LFS does not alter presynaptic vesicle density. However, we cannot rule out vesicle depletion as a potential seizure-suppressive mechanism during and immediately after LFS, as hippocampal synapses can recover from depletion within seconds (Stevens and Wesseling 1998; Abrahamsson et al. 2005).

The mechanisms of seizure control during LFS remain unclear and may differ from the mechanisms underlying longer-lasting effects. Studies with rat brain slices and 4-aminopyridine-induced epileptiform activity have suggested that the principle antiepileptic mechanism of hippocampal LFS is hyperpolarization mediated by GABA_B_ receptor activation (Toprani and Durand 2013; Smirnova et al. 2020). Accordingly, GABA_B_ receptor antagonists have been shown to diminish the antiepileptic effects of entorhinal LFS in a kindling rat model (Xu et al. 2016). The role of GABA_A_ receptors in the LFS mechanisms remains debated (Toprani and Durand 2013; Smirnova et al. 2020; Avoli et al. 2023). If LFS induces LTD in the epileptic hippocampus, the mechanism could also involve endocannabinoid release and activation of presynaptic cannabinoid-1 receptors, critical for LFS-induced LTD at Schaffer collateral synapses in CA1 (Izumi and Zorumski 2012). Thus, LFS may exert immediate antiepileptic effects through GABAergic inhibition or endocannabinoid signalling, while long-lasting effects are likely mediated by neuroplasticity, such as structural synaptic modifications.

### Hippocampal LFS reduces focal but not generalized seizures

In contrast to its effects on focal electrographic seizures, hippocampal LFS did not reduce generalized convulsive seizures in epileptic mice. We speculate that the reason is twofold: (1) generalized seizures involve larger networks beyond the seizure focus, and (2) LFS primarily modulates the activity of local hippocampal neurons. A recent functional MRI study demonstrated that 1 Hz LFS influences only the stimulated area in the sclerotic hippocampus, while higher frequencies (40-100 Hz) recruit broader brain regions, including the thalamus and neocortical areas (Schwaderlapp et al. 2024). Although MTLE is classified as focal epilepsy, functional network alterations extend beyond the sclerotic hippocampus, and seizure activity is influenced by wide-ranging networks (Spencer 2002; Bonilha et al. 2012; Bernhardt et al. 2019; Farrell et al. 2019; Vetkas et al. 2022; Padmasola et al. 2024). Retrospective studies in epilepsy patients who received DBS or RNS have shown that improved seizure control correlates with higher functional connectivity between cortical areas (Middlebrooks et al. 2018; Charlebois et al. 2022; Fan et al. 2022; Scheid et al. 2022), confirming that seizure control by neurostimulation depends on network effects.

In a previous study, we demonstrated that generalized seizures induced by optogenetic 10 Hz stimulation of PP fibers in the sclerotic hippocampus could be prevented by preconditioning with 30 min LFS (Paschen et al. 2020). Local LFS also dampened generalized seizures triggered by hippocampal kindling (Ruan et al. 2020). However, spontaneous generalized seizures may originate differently, potentially involving extrahippocampal structures or the contralateral hippocampus. In this study, contralateral hippocampal astrogliosis correlated with the occurrence of generalized seizures, supporting the idea that crosstalk between contralateral and ipsilateral hippocampi is crucial for sustained epileptic activity (Padmasola et al. 2024). Therefore, local hippocampal LFS alone may be insufficient to suppress generalized seizures, and broader network engagement may be necessary either via neurostimulation or pharmacological treatment as generalized seizures are commonly drug-responsive (Klein et al. 2015; Duveau et al. 2016)

### Hippocampal LFS ameliorates spatial memory deficits in epileptic mice

The hippocampus is central for spatial navigation and memory (Burgess et al. 2002; Buzsáki and Moser 2013), functions often compromised in patients and rodent models of MTLE (Gröticke et al. 2008; Bell et al. 2011; Tramoni-Negre et al. 2017; Van Den Herrewegen et al. 2018; Paschen et al. 2024). In the present study, epileptic mice showed long-term spatial memory impairments which were slightly alleviated by prolonged hippocampal LFS. More specifically, KA mice made more errors and required more time to reach the target in the Barnes maze test than healthy controls, whereas the performance of KA+LFS mice was comparable to controls. Consistent with previous work (Van Den Herrewegen et al. 2018; Paschen et al. 2024), epileptic mice relied on a non-spatial serial search strategy, unlike healthy controls, which used spatial cues, went to the target directly, and had a clear spatial preference for the target quadrant. The enhanced fear of being in the center or thigmotaxis, found in epileptic mice in the open-field test, might have contributed to the selection of serial search strategy. In contrast to KA mice, KA+LFS mice spent more than 25% of the time in the target quadrant, showed a greater tendency to use a spatial search strategy, and were less likely to fail in reaching the target. These findings suggest that long-term hippocampal LFS provides partial improvement but does not fully restore spatial memory in MTLE with HS.

Hippocampal LFS has been shown to restore spatial learning and memory in the pilocarpine-induced rat model of epilepsy (Wang et al. 2020; Zare et al. 2024), which might be feasible in a model with mild hippocampal sclerosis. Although the extent of hippocampal sclerosis does not directly correlate with spatial memory (Glikmann-Johnston et al. 2008; Paschen et al. 2024), hippocampal damage most likely contributes to memory impairment. Our data suggest that while LFS does not alter hippocampal sclerosis, it may alleviate memory deficits through seizure suppression or stimulation-induced synaptic changes. When a dentate gyrus network model was modulated to have increased PP-DGC input strength, the pattern separation ability of the dentate gyrus was compromised (Yim et al. 2015), suggesting that the LTP-like ultrastructural changes might contribute to memory impairment, which in turn can be improved by reversal of these alterations by LFS.

### Limitations and perspectives

This study has some limitations and poses questions for future investigation. First, we terminated our *in vivo* experiments immediately after the last LFS session, enabling us to assess histopathology, neurogenesis, and ultrastructure in the time frame during which we expected LFS-induced effects based on our previous studies(Paschen et al. 2020; Paschen et al. 2024). Future research focusing on the time course of the LFS effects should include electrophysiological and ultrastructural analysis after extended periods. Second, we did not monitor the oestrous cycle in female mice, which is known to influence seizure activity (Li et al. 2020). Some studies suggest this only applies to mice with KA injections in the left hippocampus (Cutia et al. 2023), but it remains an important consideration.

In light of clinical significance, it is crucial to consider how well our findings in epileptic mice translate to humans. Although the intrahippocampal KA mouse model is well-suited for MTLE with hippocampal sclerosis, there are relevant differences between humans and mice, such as seizure frequency, clinical manifestations associated with seizures, diurnality, and brain complexity. In MTLE patients, focal seizures are associated with olfactory and gustatory auras, visceral sensations, feelings of déjà vu, and confusion, which cannot be analysed in mice. Other clinical features of focal seizures include oral and manual automatisms and behavioural arrest (Maillard et al. 2004; Fisher et al. 2017; Vinti et al. 2021). These subtle changes are complicated to detect in mice by simple video observation, which might be improved by machine-learning-assisted video analysis in the future (Gschwind et al. 2023). Another aspect is that mice are nocturnal, and we stimulated them during the light phase translating to night-time stimulation in humans. Diurnal seizure cycles are patient-specific (Karoly et al. 2016; Leguia et al. 2021; Bernard et al. 2023), thus the stimulation time should be individually fine-tuned. The LFS parameters, especially injected current, need to be adjusted in a larger brain, which has already been successfully implemented (Velasco et al. 2007; Cukiert et al. 2017; Nair et al. 2020). We found no adverse effects of LFS on cognition or histopathological parameters, encouraging further assessment of LFS in controlled clinical trials.

## Conclusion

In summary, long-term hippocampal LFS reliably suppressed focal seizures in chronically epileptic female and male mice, with seizure reduction beyond the stimulation period. LFS also improved spatial memory and reversed pathological changes at PP-DGC synapses but did not affect generalized seizures, neurogenesis, and hippocampal sclerosis. These findings suggest that hippocampal LFS could be a promising therapeutic approach for MTLE patients that are non-responders to anti-seizure medications and HFS.

## Abbreviations

AAV: adeno-associated virus
CA: Cornu ammonis
CaMKIIα: Ca^2+/^calmodulin-dependent protein kinase II α
CEMT: Center for Experimental Models and Transgenic Service
cHC: contralateral hippocampus
DBS: deep brain stimulation
DGC: dentate granule cell
Doublecortin: DCX
GFAP: glial fibrillary acidic protein
HFS: high-frequency stimulation
HL: high-load
iHC: ipsilateral hippocampus
KA: kainate
LFP: local field potential
LFS: low-frequency stimulation
LL: low-load
LTD: long-term depression
LTP: long-term potentiation
ML: medium load
MML: middle molecular layer
MTLE: mesial temporal lobe epilepsy
NeuN: neuronal nuclei
PB: phosphate buffer
PFA: paraformaldehyde
PP: perforant path
PSD: postsynaptic density
RM: repeated measures
RNS: responsive neurostimulation
ROI: region of interest
Sal: Saline
vGluT1: vesicular glutamate transporter 1

## Acknowledgments

We thank Christoph Janus and Alejandro Perez Recio for their support with data analysis. We are very grateful for the technical assistance by Sigrun Nestel from the Department of Neuroanatomy. We thank Dr Christian Böhler, formerly Bioelectronic Microtechnology - IMTEK, for providing laboratory equipment and supervision for electrode coatings.

This work was supported by the German Research Foundation grant HA 1443/12-1.

## Author contributions

**PK**: Conceptualization; Data curation; Formal analysis; Investigation; Project administration; Visualization; Writing - original draft; **EP**: Conceptualization; Data curation; Methodology; **ADM**: Investigation; Formal analysis; **AV**: Investigation; Resources; Writing – review & editing; **CAH**: Conceptualization; Funding acquisition; Project administration; Resources; Supervision; Writing - review & editing; **UH**: Conceptualization; Project administration; Methodology; Validation; Writing – review & editing;

