## Supplementary Figures 1-6, Supplementary Table 1 for "Long-term hippocampal low-frequency stimulation alleviates focal seizures, memory deficits and synaptic pathology in epileptic mice"

### Inventory of Supplemental Information

Figure S1, Related to Methods

Figure S2, Related to Figure 1

Figure S3, Related to Figure 2

Figure S4, Related to Figure 3

Figure S5, Related to Figure 4

Figure S6, Related to Figures 5

Supplemetary Table, related to Methods

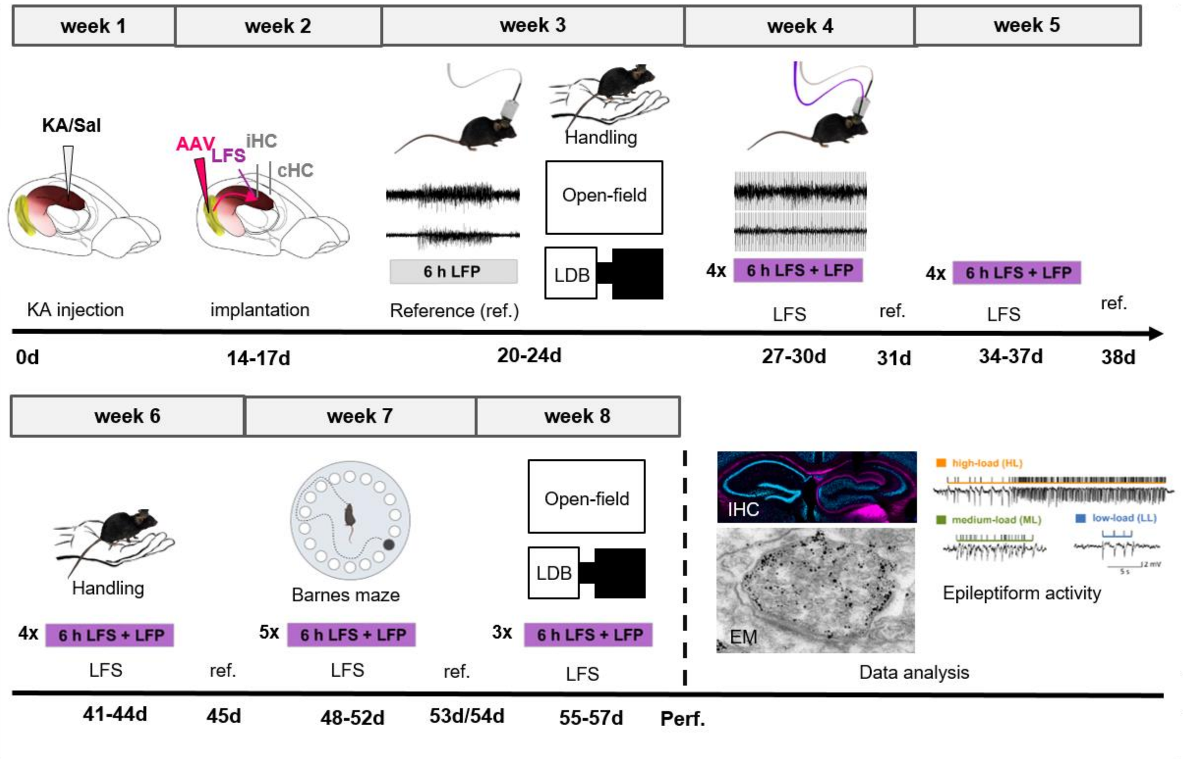


**Figure S1. Timeline of experiments.** Animals received intrahippocampal KA or Sal into the right dorsal hippocampus. After 2 weeks, the mice were implanted with recording electrodes in both hippocampi (iHC, cHC) and a stimulation electrode for LFS next to the iHC electrode. A viral vector (AAV) carrying mCherry as fluorescent reporter under control of the CamKII promotor was injected into the medial entorhinal cortex to label perforant path fibers in a subset of mice that later underwent electron microscopy (EM). 6-h local field potential (LFP) recordings were performed without LFS (reference - ref) once a week and with LFS four times a week for one month. Mice were subjected to handling and the following behavioral tests: open-field, light-dark box (LDB), and Barnes maze. Transcardial perfusion (perf.) was performed on day 57 after intrahippocampal injections and the tissue was processed for immunohistochemistry (IHC) and EM in a subset of mice.

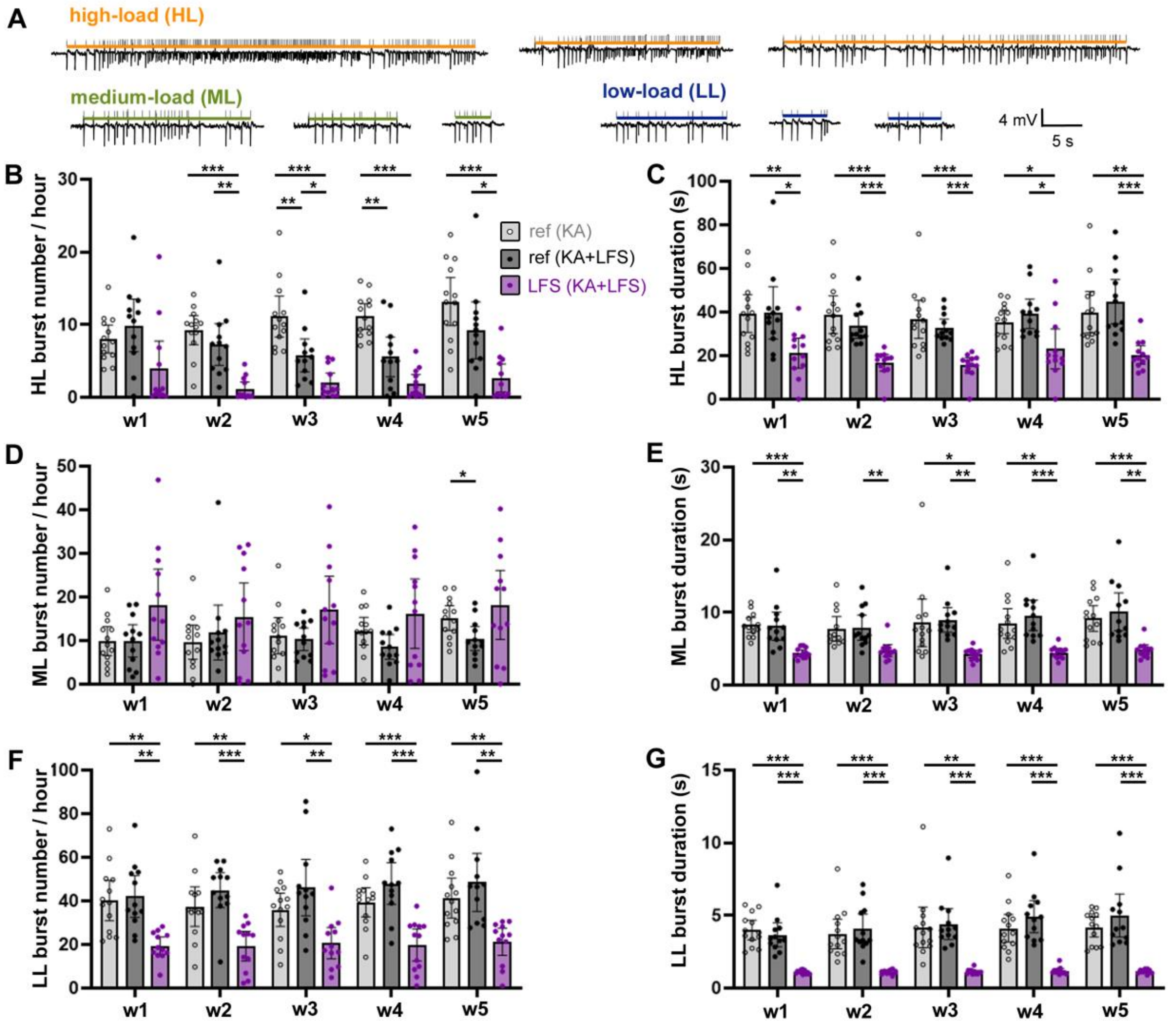


**Figure S2. The effects of long-term LFS on epileptiform burst number and duration**. (A) Examples of high-load (HL), medium-load (ML), and low-load (LL) bursts classified with the PEACOC algorithm. (B) HL burst numbers (per hour) in reference (ref) recordings from KA mice (grey, n=13) and KA+LFS mice (dark grey, n=12) and during LFS (purple) across five stimulation weeks. Two-way RM ANOVA (interaction: $F_{(8,\ 136)}$=2.80, P=0.0067, time factor: $F_{(2{,}804,\ 95{,}33)}$=2.91, P=0.04, group factor: $F_{(2,\ 34)}$=31.09, P<0.0001). (C) HL burst durations. Two-way RM ANOVA (interaction: $F_{(8,\ 136)}$=0.96, P=0.47, time factor:, $F_{(3{,}481,\ 118{,}4)}$=2.79, P=0.036, group factor: $F_{(2,\ 34)}$=19.72, P<0.0001). (D) ML burst numbers, Two-way RM ANOVA (interaction: $F_{(8,\ 136)}$=0.83, P=0.57, time factor: $F_{(2{,}774,\ 94{,}33)}$=0.87, P=0.45, group factor: $F_{(2,\ 34)}$=3.61, P=0.038). (E) ML burst durations. Two-way RM ANOVA (interaction: $F_{(8,\ 136)}$=0.74, P=0.66, time factor: $F_{(3{,}046,\ 103{,}5)}$=2.38, P=0.073, group factor: $F_{(2,\ 34)}$=15.58, P<0.0001). (F) LL burst numbers. Two-way RM ANOVA (interaction: $F_{(8,\ 136)}$=0.26, P=0.98, time factor: $F_{(2{,}698,\ 91{,}72)}$=0.47, P=0.68, group factor: $F_{(2,\ 34)}$= 28.44, P<0.0001). (G) LL burst durations. Two-way RM ANOVA (interaction: F(8, 136)=0.92, P=0.50, time factor: $F_{(2{,}917,\ 99{,}18)}$=1.87, P=0.14, group factor: $F_{(2,\ 34)}$=38.94, P<0.0001). Tukey's multiple comparisons test: *P<0.05, **P<0.01, ***P<0.001. Data are presented as mean with 95% CI.

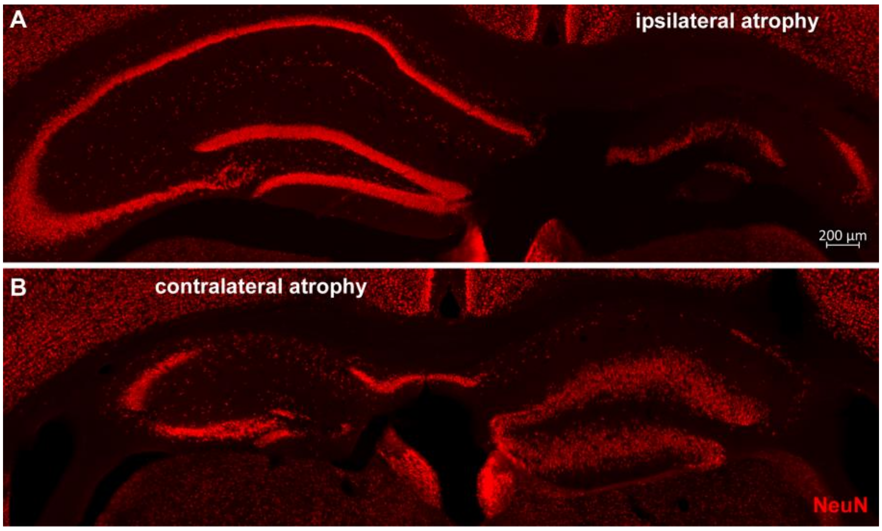


**Figure S3. Examples of severe hippocampal atrophy.** (A) NeuN-labeling showing a strong atrophy in the ipsilateral hippocampus, including complete loss of CA1, CA3 and hilar neurons and unusual loss of dentate granule cells. (B) Abnormal atrophy of contralateral hippocampus with neuronal loss in the CA1 and dentate gyrus.

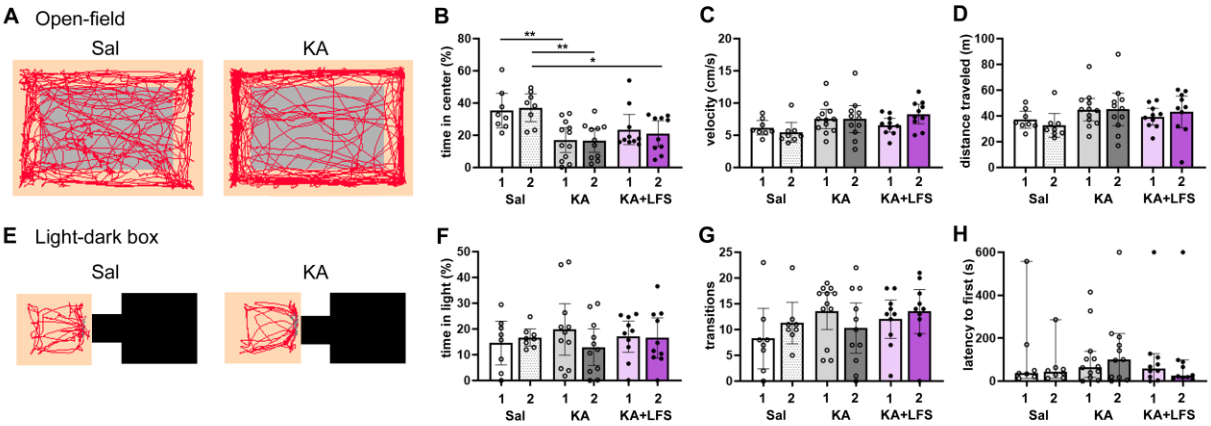


**Figure S4. Mobility and anxiety-like behavior in open-field and light-dark box tests.** (A) Movement trajectory examples from Sal and KA mice in the open-field test (10 min). The center of the arena is shown in grey. (B) Percentage of time spent in the center of the open-field arena by Sal, KA, and KA+LFS mice 22 days (1) and 56 days post-KA/Sal injection (2), corresponding to before and after LFS for KA-LFS mice. Mixed effects analysis (interaction: $F_{(2, 26)}$=0.17, P=0.84, time factor: $F_{(1, 26)}$=0.011, P=0.92, group factor: $F_{(2, 27)}$=12.10, P=0.0002), Šídák's multiple comparisons test. (C) Velocity (cm/s) in the open-field test. Mixed effects analysis (interaction: $F_{(2, 26)}$= 1.72 P=0.20, time factor: $F_{(1, 26)}$=0.35, P=0.56, group factor: $F_{(2, 27)}$= 3.05, P= 0.064). (D) Distance traveled (m) in open-field test. Mixed effects analysis (interaction: $F_{(2, 53)}$= 0.36, P=0.70, time factor: $F_{(1, 53)}$= 0.0019, P= 0.96, group factor: $F_{(2, 53)}$=2.46 P=0.10). (E) Examples of trajectories made by Sal and KA mice in the light compartment (orange) in the light-dark box test (10 min). (F) Fraction of time spent in the light compartment. Mixed effects analysis (interaction: $F_{(2, 26)}$= 0.28, P=0.75, time factor: $F_{(1, 26)}$= 0.035, P=0.85, group factor: $F_{(2, 27)}$=0.26, P=0.77). (G) Number of transitions from dark to light compartment. Mixed effects analysis (interaction: $F_{(2, 26)}$= 2.93, P=0.07, time factor: $F_{(1, 26)}$= 0.13, P=0.73, group factor: $F_{(2, 27)}$=0.77, P= 0.47). (H) Latency to the first transition (if 600s, then mouse did not come out). Kruskal-Wallis test (P=0.88). *P<0.05, **P<0.01. Data are presented as mean (B-D, F-G) or median (H) with 95% CI.

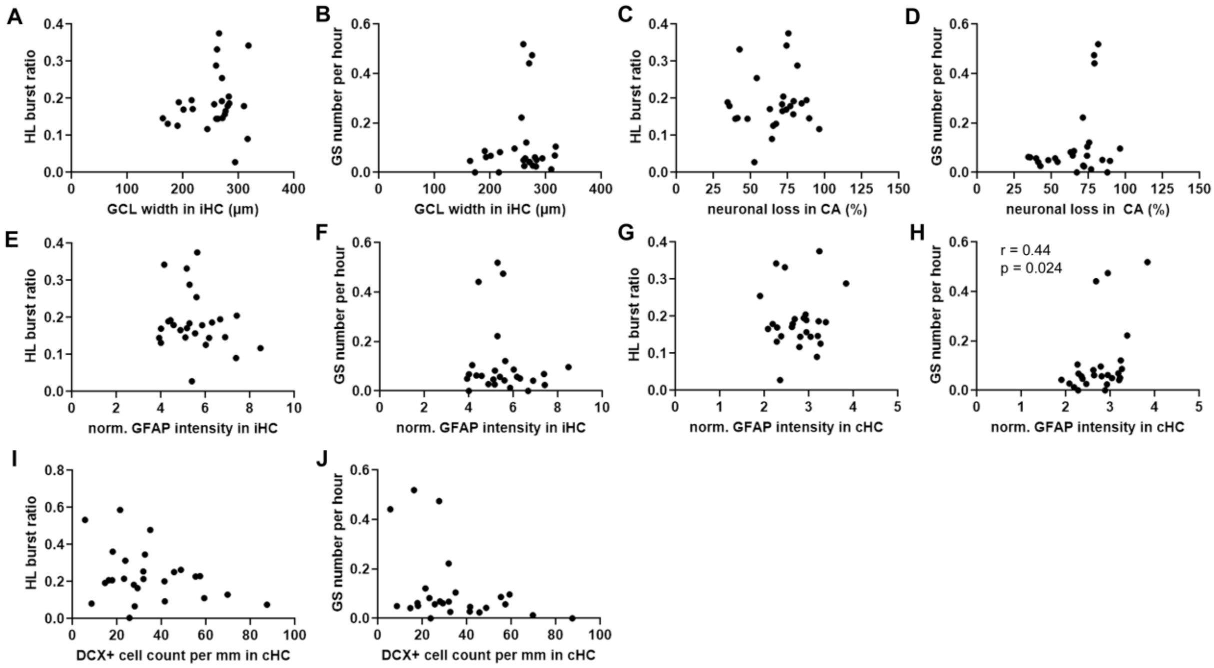


**Figure S5. Correlations between histological findings and seizure activity.** Relationship between: (A) granule cell layer (GCL) width in ipsilateral hippocampus (iHC) and average HL burst ratio in reference recordings. (B) granule cell layer (GCL) width in iHC and average generalized seizure (GS) number per hour. (C, D), neuronal loss in CA area with (C) HL burst ratio / (D) generalized seizure number. (E, F) normalized GFAP intensity in iHC and (E) HL burst ratio / (F) generalized seizure number. (G, H), normalized GFAP intensity in contralateral hippocampus (cHC) and HL burst ratio (G) / generalized seizures (H). (I, J) DCX-positive cell count in cHC and (I) HL burst ratio / (J) generalized seizures. Pearson's correlation test, n=26.

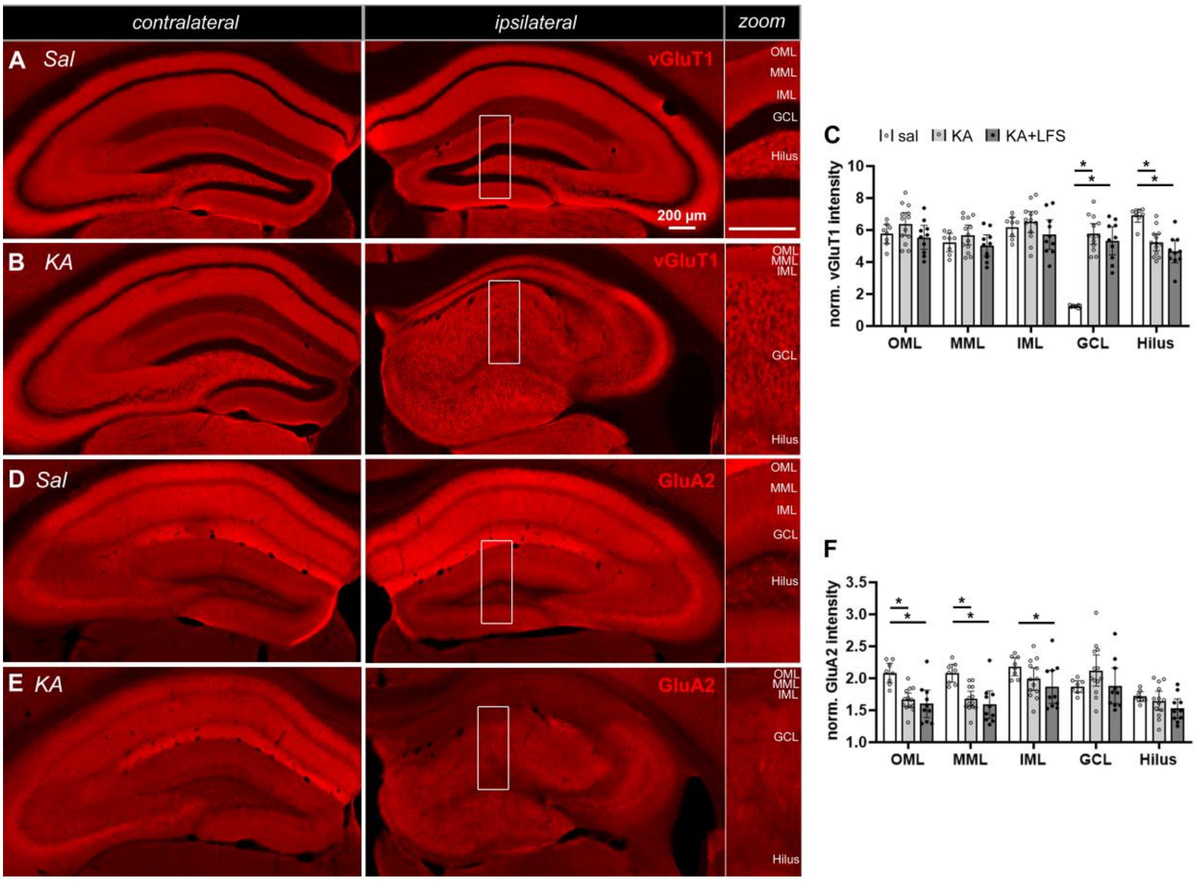


**Figure S6. Presynaptic and postsynaptic markers for excitatory synapses revealed KA-induced, but not LFS-induced, changes in the dentate gyrus.** (A, B) Hippocampal sections with vGluT1 labeling excitatory presynaptic terminals in (A) healthy and (B) epileptic mice. Scale bars 200 µm. OML- outer molecular layer, MML- middle molecular layer, IML- inner molecular layer, GCL-granule cell layer. (C) Normalized vGluT1 intensity in the five layers of the dentate gyrus. Two-way ANOVA (interaction: $F_{(8, 140)}$=16.53, P<0.0001, layer: $F_{(4, 140)}$=9.97, P<0.0001, group factor: $F_{(2, 140)}$=13.45, P<0.0001), Tukey's multiple comparisons tests (for group factor). (D, E) Hippocampal sections with GluA2 staining for the type2 AMPA receptor subunits in the excitatory dendrites. (F) Normalized GluA2 intensity in the five layers of the dentate gyrus. Two-way ANOVA (interaction: $F_{(8, 140)}$=2.81, P=0.0064, layer: $F_{(4, 140)}$=9.97, P<0.0001, group factor: $F_{(2, 140)}$=13.45, P<0.0001), Tukey's multiple comparisons tests (for group factor). *P<0.05. Data are presented as mean with 95% CI.

**Table S1. Overview of the sample sizes and excluded animals, categorized by exclusion criteria.** *One animal lost the LFP signal 30 days after KA but was included in histology and behavioral analysis. **One animal was excluded from focal seizure analysis due to elevated GS in the last recording weeks, which hindered adequate assessment of the HL burst ratio. ***Mice that were frozen or jumped off the Barnes Maze (BM) arena.

| Group | Females (Sal) | Males (Sal) | Females (KA) | Males (KA) |
|---|---|---|---|---|
| total injected | 4 | 6 | 30 | 36 |
| died during surgery | 0 | 0 | 3 | 3 |
| died during/after SE (<4 days) | - | - | 4 | 8 |
| sudden death (>4 days after injection) | 0 | 1 | 1 | 3 |
| insufficient recovery from surgery | 1 | 0 | 1 | 1 |
| lack of SE, HS or recurrent seizures | - | - | 4 | 2 |
| implant came off | - | - | 3 | 1 |
| abnormal hippocampal atrophy | - | - | 2 | 4 |
| sample size (LFP, histology) | 3 | 5 | 12 | 14 |
| sample size: non-stimulated | | | 6/7* | 7 |
| sample size: stimulated (KA+LFS) | | | 5/6** | 7 |
| abnormal behavior in BM*** | 0 | 0 | 4 | 0 |
| sample size (behavior) | 8 | | 22 | |
| sample size: non-stimulated | | | 12 | |
| sample size: stimulated (KA+LFS) | | | 10 | |
| sample size (electron microscopy) | | | | 8 |
| no immunogold labeling | | | | 2 |
| sample size: non-stimulated | | | | 3 |
| sample size: stimulated (KA+LFS) | | | | 3 |
